# Statistical notes on false positive and negative error rates in the evaluation of long-term carcinogenicity bioassays

**DOI:** 10.1101/2022.02.25.481968

**Authors:** Ludwig A. Hothorn

## Abstract

The appropriate interpretation of mortality-adjusted tumor incidences in long-term carcinogenicity bioassays depends substantially on the actual false positive and false negative error rates. These depend, among other things, on the type of analysis of multiple correlated tumor sites and the mode of dose-response dependence in relation to the design. Selected quantitative results, such as shape-to-design relationship and discreteness are presented and the influence of further issues is discussed qualitatively.

## 1 The problem

Recently, a discussion on the evaluation of mortality-adjusted tumor rates for multiple correlated tumors using min-p poly-3 trend tests was published [7]. A rejoinder [31] characterized this approach as unacceptably conservative from a toxicological perspective (this rejoinder was answered by the first authors [8]). In essence, the first paper considers the multiplicity-adjusted analysis of multiple tumors (and their additional further summaries) using min-p test, a permutation-based maximum-type union-intersection test (UIT) over the *q* tumors (respectively *q* + *q*^*′*^ summaries). The re-jointing paper lists several counter-arguments from a statistical point of view besides inadequacy of p-values, the use of historical controls (especially for rare tumors) and especially the increased conservatism of the multiplicity correction considering multiple correlated tumors. The following historical quote is central to these arguments: *‘Finally, it has been recognized that a primary difficulty with animal bioassay data is not high false positive rates, but low statistical power, and that false positives are primarily an issue with common tumors (Portier, 1994)’* [28]. I.e. the essential and for the following statistical properties central issue is the ratio of false positive to false negative error rate. This ratio is quite complex and therefore several statistical aspects are considered in detail, particularly when considering multiplicity.

In view of the relevance of the multiplicity problem, it is disillusioning that the only clear definition of the problem is to be found in an FDA guidance that is more than 20 years old (still in draft status!) [1] *‘Because of the large number of comparisons involved (usually 2 species, 2 sexes, and 30 or more tissues examined), a great potential exists for finding statistically significant positive trends or treatment-placebo differences due to chance alone (i*.*e*., *a false positive). Therefore, it is important that an overall evaluation of the carcinogenic potential of a drug take into account the multiplicity of statistical tests of significance for both trends and pairwise comparisons’*.

First, we discuss the *f*^+^*/f*^−^ association for the independent analysis multiple tumor sites compared to joint analysis. Furthermore, this relationship is considered for sources and multiplicity: i) multiple studies, ii) trend and pairwise tests, iii) multiple effect sizes, iv) multiple Weibull shape parameter k of poly-k tests, and v) multiple trend alternatives. Furthermore, this correlation is quantified for the underlying Armitage trend tests in the unbalanced design for different shapes and *p*_0_. Furthermore, the exhausting the size of the test statistics with respect of the serious discreteness of the 2 by k table data is considered.

## 2 False positive vs. false negative error rates in the evaluation of multiple tumors

Commonly *>* 50 tumor sites are diagnosed in these bioassays, those that have been defined a priori and others that occur spontaneously. In addition, there are content summaries, such as (adenomas of the lymphatic system). Analyzing these primary endpoints independently, each at the *α* level, the false positive rate of the experiment increases massively towards 1. If these primary correlated endpoints are analyzed together, and a procedure is available for the poly-k test [12], the false negative rate increases massively (with compliance to the FWER). This increase is more pronounced the more and less correlated tumors are considered. A joint evaluation of all tumors is technically possible, but appears to be unacceptably conservative. The consideration of the discreteness of these rare events leading to the minimization of the *f*^−^ rate is available for crude proportions (and for pairwise comparisons only), but probably not for the poly-k test [12].

The fundamental dilemma of both approaches: independent analysis: *f*^+^ ⇒ 1, but joint analysis: *f*^−^ ⇒ 1 is not easily resolvable. One could avoid too high *f*^−^ rates in the joint approach by not taking into account the numerous tumor sites with very small incidences as well as its independent application to body systems only [5]. As long as there is no consensus, preferably within a guideline, it is of little help to reject the other approach as indisputable. What would I do as a biostatistician in the meantime: evaluate the data with both approaches simultaneously, and critically discuss the few, really relevant, findings with the toxicologists. No case like this, *p*_*xyz*_ *<* 0.05, and thus the substance is carcinogenic. The statistical methods and the software for this are available, with the restriction of suitable consideration of the discreteness for the exhaustion of the test level.

What could be a way out of this dilemma of the two extreme approaches? A compromise could be the FWER control within body systems, like e.g. respiratory, lymphoreticular, etc. after [5], but per organ system testing to the level *α*.

## 3 False positive vs. false negative rates for the Armitage trend test as the basis of the poly-k test

The poly-k test is mentioned in a recent recommendation [24] and currently appears to be the standard of evaluation of mortality-adjusted tumor rates, see e.g. [38]. The question arises as to what the ratio *f*^+^*/f*^−^ for the Armitage trend test is as the basis for the poly-k test.

### 3.1 False negative rates for various dose-response shapes and several spontaneous rates *p*_0_’s

The impact of several shapes and *p*_0_’s on Armitage trend tests *f*^−^ rate is characterized using an approximation in the CRAN package *multiCA* [34] ^1^. Table 1 shows a superiority ratio 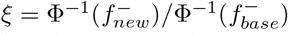 relative to the base scenario (i.e. balanced NTP design *n*_*i*_ = 50, 50, 50, 50), where the greater *ξ*, the more superior is the particular condition:

**Table 1:** Impact of various balanced and unbalanced designs on superiority parameter *ξ*

| No. | Proportions | Samples sizes | Dose levels | f- | Scenario | ξ-factor |
| --- | --- | --- | --- | --- | --- | --- |
| 1 | 0.01,0.05,0.15,0.2 | 50,50,50,50 | 0,10,50,100 | 0.0702 | base $N^{NTP}$ | - |
| 2 | 0.01,0.05,0.15,0.2 | 40,40,40,40 | 0,10,50,100 | 0.1331 | reduced balanced $n_i$ | 75 |
| 3 | 0.01,0.05,0.15,0.2 | 30,30,30,30 | 0,10,50,100 | 0.2420 | strongly reduced balanced $n_i$ | 48 |
| 4 | 0.01,0.05,0.15,0.2 | 60,60,60,60 | 0,10,50,100 | 0.0361 | increased balanced $n_i$ | 122 |
| 5 | 0.01,0.05,0.15,0.2 | 100,50,50,50 | 0,10,50,100 | 0.0099 | dual control unbalanced $n_i$ | 158 |
| 6 | 0.01,0.05,0.15,0.2 | 80,40,40,40 | 0,10,50,100 | 0.0302 | dual control unbalanced (N=const) | 127 |
| 7 | 0.01,0.05,0.15,0.2 | 74,42,42,42 | 0,10,50,100 | 0.0355 | $n_0 = \sqrt{(n)}$ unbalanced (N=const) | 122 |
| 8 | 0.01,0.05,0.15,0.2 | 86,38,38,38 | 0,10,50,100 | 0.0257 | unbalanced (N=const) | 132 |
| 9 | 0.01,0.05,0.15,0.2 | 89,37,37,37 | 0,10,50,100 | 0.0238 | unbalanced (N=const) | 134 |
| 10 | 0.01,0.05,0.15,0.2 | 92,36,36,36 | 0,10,50,100 | 0.0221 | unbalanced (N=const) | 137 |
| 11 | 0.01,0.05,0.15,0.2 | 95,35,35,35 | 0,10,50,100 | 0.0204 | unbalanced (N=const) | 139 |
| 12 | 0.01,0.05,0.15,0.2 | 98,34,34,34 | 0,10,50,100 | 0.0190 | unbalanced (N=const) | 141 |
| 13 | 0.01,0.05,0.15,0.2 | 101,33,33,33 | 0,10,50,100 | 0.0177 | unbalanced (N=const) | 143 |
| 14 | 0.01,0.05,0.15,0.2 | 104,32,32,32 | 0,10,50,100 | 0.0165 | unbalanced (N=const) | 145 |
| 15 | 0.01,0.05,0.15,0.2 | 107,31,31,31 | 0,10,50,100 | 0.0155 | unbalanced (N=const) | 146 |
| 16 | 0.01,0.05,0.15,0.2 | 110,30,30,30 | 0,10,50,100 | 0.0145 | unbalanced (N=const) | 148 |
| 17 | 0.01,0.05,0.15,0.2 | 125,25,25,25 | 0,10,50,100 | 0.0109 | unbalanced (N=const) | 156 |
| 18 | 0.01,0.05,0.15,0.2 | 140,20,20,20 | 0,10,50,100 | 0.0092 | unbalanced (N=const) | 160 |
| 19 | 0.01,0.05,0.15,0.2 | 155,15,15,15 | 0,10,50,100 | 0.0092 | unbalanced (N=const) | 160 |
| 20 | 0.01,0.05,0.15,0.2 | 170,10,10,10 | 0,10,50,100 | 0.0130 | unbalanced (N=const) | 151 |
| 21 | 0.01,0.05,0.15,0.2 | 185,10,10,10 | 0,10,50,100 | 0.0402 | unbalanced (N=const) | 119 |
| 22 | 0.01,0.05,0.15,0.2 | 20,20,20,140 | 0,10,50,100 | 0.274 | inverse unbalanced (N=const) | 41 |
| 23 | 0.01,0.05,0.15,0.2 | 50,50,45,40 | 0,10,50,100 | 0.081 | unbalanced $n_i$ due to mortality $N < N^{NTP}$ | 95 |
| 24 | 0.01,0.05,0.15,0.2 | 65,50,45,40 | 0,10,50,100 | 0.047 | unbalanced $n_i$ due to mortality (N=const) | 114 |
| 25 | 0.01,0.05,0.15,0.2 | 47,46,46,46 | 0,10,50,100 | 0.0877 | balanced $n_i$ due to mortality $N < N^{NTP}$ | 92 |
| 27 | 0.01,0.05,0.15,0.2 | 50,45,30,25 | 0,10,50,100 | 0.116 | strong unbalanced $n_i$ due to mortality $N < N^{NTP}$ | 81 |
| 28 | 0.01,0.05,0.15,0.2 | 100,45,30,25 | 0,10,50,100 | 0.0231 | strong unbalanced $n_i$ due to mortality N=const | 135 |
| 29 | 0.01,0.05,0.15,0.2 | 39,37,37,37 | 0,10,50,100 | 0.149 | balanced $n_i$ due to mortality $N < N^{NTP}$ | 71 |

First in Table 1 reveals the old familiar relationship that *f*^−^ decreases with increasing *n*_*i*_ (vice versa) (see Scenarios 2,3,4). No. 5 shows the massive decrease of the *f*^−^ rate (and thus increase of *ξ*) for the sometimes used dual control design, mainly due to the higher N. A first simple conclusion for toxicologists arises from comparison reasons of *f*^−^ to use the NTP design (*n*_*i*_ = 0, 50, 50, 50) by itself, even if it is not power-optimal (see below). For a fair comparison the impact of partially unbalanced design is demonstrated for constant total sample size *N* in the scenarios 6-21 whereas *f*^−^ decreases with higher *n*_0_ *> n*_*i*_ up to irrelevant high unbalancedness e.g. in case 19. Simply, *f*^−^ decreases the more smaller *p*_*i*_ occur in larger *n*_*i*_, although the pattern of the *p*_*i*_’s is not known a-priori. From a statistical point of view of a minimal *f*^−^ rate, this results in a degree of unbalance much greater than dual control design, which may be difficult to communicate to toxicologists. Another conclusion seems to be the direct inclusion of historical controls (see below). Not surprisingly, inverse unbalancing leads to a collapse of *ξ*. Partially unbalanced designs due to mortality-related lower *n*_*i*_’s decreases *ξ* due to smaller *N* but does not lead to a more pronounced increase of *f*^−^, especially when compared to the unrealistic N=const unbalanced design (no. 23-29). The third encouraging conclusion for toxicologists is the relative small increase in *f*^−^ rate with mortality-related reduced *n*_*i*_ in the dose groups.

Second, Table 2 reveals the association between shape of the dose-response curve (near-to-linear no. 3-4, plateau-shape no. 5-8 and sub-linear no. 9-12) and selected dose scores. One reason for the *f*^−^ increase in plateau shapes is the increase in global *π* and thus the common variance *π*(1 − *pi*). The increase of *f*^−^ can be massive with inappropriately chosen scores - also an indication not to use the standard test for the linear trend, but e.g. the Tukey trend test [32].

**Table 2:** Impact of dose scores on superiority parameter *ξ*

| No. | Proportions | Samples sizes | Dose levels | f- | Scenario | ξ-factor |
| --- | --- | --- | --- | --- | --- | --- |
| 1 | 0.01,0.05,0.15,0.2 | 50,50,50,50 | 0,10,50,100 | 0.0702 | base $N^{\text{NTP}}$ | - |
| 2 | 0.01,0.05,0.15,0.2 | 50,50,50,50 | 0,1,2,3 | 0.0627 | ordinal dose scores | 104 |
| 3 | 0.01,0.05,0.15,0.2 | 50,50,50,50 | 0,10,11,12 | 0.193 | plateau-style dose scores | 59 |
| 4 | 0.01,0.05,0.15,0.2 | 50,50,50,50 | 0,10,20,40 | 0.0737 | lin-log dose scores | 98 |
| 5 | 0.01,0.2,0.2,0.2 | 50,50,50,50 | 0,10,50,100 | 0.524 | base dose scores | (< 1) |
| 6 | 0.01,0.2,0.2,0.2 | 50,50,50,50 | 0,1,2,3 | 0.293 | ordinal dose scores | 37 |
| 7 | 0.01,0.2,0.2,0.2 | 50,50,50,50 | 0,10,11,12 | 0.107 | plateau-style dose scores | 84 |
| 8 | 0.01,0.2,0.2,0.2 | 50,50,50,50 | 0,10,20,40 | 0.401 | lin-log dose scores | 17 |
| 9 | 0.01,0.01,0.01,0.2 | 50,50,50,50 | 0,10,50,100 | 0.0074 | base dose scores | 165 |
| 10 | 0.01,0.01,0.01,0.2 | 50,50,50,50 | 0,1,2,3 | 0.0280 | ordinal dose scores | 130 |
| 11 | 0.01,0.01,0.01,0.2 | 50,50,50,50 | 0,10,11,12 | 0.387 | plateau-style dose scores | 19 |
| 12 | 0.01,0.01,0.01,0.2 | 50,50,50,50 | 0,10,20,40 | 0.0076 | lin-log dose scores | 165 |

Third, Table 3 reveals the association between the shape of the dose-response curve and the spontaneous tumor rate *p*_0_. The well known massive increase of *f*^−^ with increasing *p*_0_ as a function of the variance *π*(1 − *π*) (where *π* is the common variance) is demonstrated in cases 2-5. This also applies, for the somewhat fairer comparison for Δ*p* = *const* in no. 6-9. The important conclusion for toxicologists is the necessary classification into rare, i.e. *p*_0_ *<<* 0.05 and common tumors, i.e. *p*_0_ *>* 0.05. The most massive effect, however, is in the shape of dose-response dependence: there is a significant increase in *ξ* for a sub-linear profile (compared to near-to-linear), but on the other hand a catastrophic decrease for plateau-like profiles or even downturn at high dose. Since the profile is not an assumption but a result of the study and all possible shapes can occur in different tumors even within one study, a trend test for linear shapes is contra-indicated, but e.g. the Tukey trend test [32].

**Table 3:**
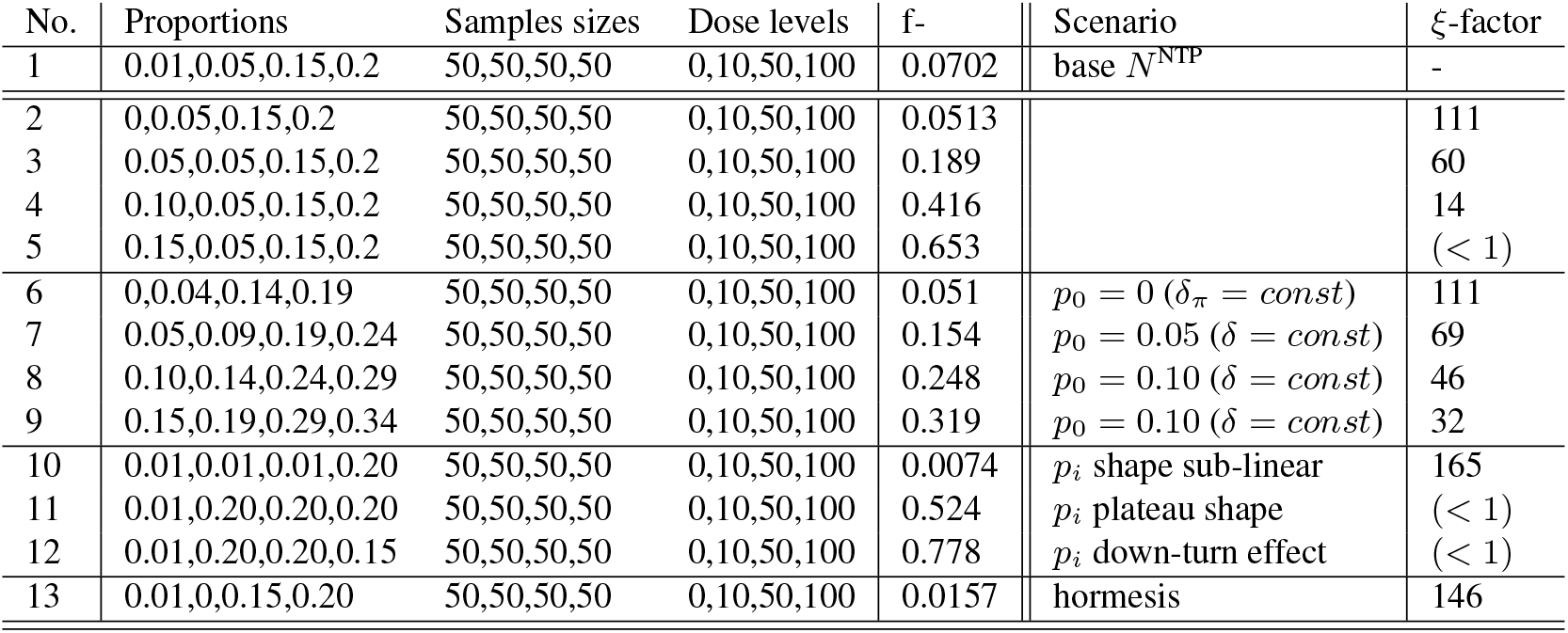
Impact of dose-response shapes and *p*_0_’s on superiority parameter *ξ*

| No. | Proportions | Samples sizes | Dose levels | f- | Scenario | ξ-factor |
| --- | --- | --- | --- | --- | --- | --- |
| 1 | 0.01,0.05,0.15,0.2 | 50,50,50,50 | 0,10,50,100 | 0.0702 | base $N^{\text{NTP}}$ | - |
| 2 | 0,0.05,0.15,0.2 | 50,50,50,50 | 0,10,50,100 | 0.0513 |  | 111 |
| 3 | 0.05,0.05,0.15,0.2 | 50,50,50,50 | 0,10,50,100 | 0.189 |  | 60 |
| 4 | 0.10,0.05,0.15,0.2 | 50,50,50,50 | 0,10,50,100 | 0.416 |  | 14 |
| 5 | 0.15,0.05,0.15,0.2 | 50,50,50,50 | 0,10,50,100 | 0.653 |  | (< 1) |
| 6 | 0,0.04,0.14,0.19 | 50,50,50,50 | 0,10,50,100 | 0.051 | $p_0 = 0$ ($\delta_\pi = \text{const}$) | 111 |
| 7 | 0.05,0.09,0.19,0.24 | 50,50,50,50 | 0,10,50,100 | 0.154 | $p_0 = 0.05$ ($\delta = \text{const}$) | 69 |
| 8 | 0.10,0.14,0.24,0.29 | 50,50,50,50 | 0,10,50,100 | 0.248 | $p_0 = 0.10$ ($\delta = \text{const}$) | 46 |
| 9 | 0.15,0.19,0.29,0.34 | 50,50,50,50 | 0,10,50,100 | 0.319 | $p_0 = 0.10$ ($\delta = \text{const}$) | 32 |
| 10 | 0.01,0.01,0.01,0.20 | 50,50,50,50 | 0,10,50,100 | 0.0074 | $p_i$ shape sub-linear | 165 |
| 11 | 0.01,0.20,0.20,0.20 | 50,50,50,50 | 0,10,50,100 | 0.524 | $p_i$ plateau shape | (< 1) |
| 12 | 0.01,0.20,0.20,0.15 | 50,50,50,50 | 0,10,50,100 | 0.778 | $p_i$ down-turn effect | (< 1) |
| 13 | 0.01,0,0.15,0.20 | 50,50,50,50 | 0,10,50,100 | 0.0157 | hormesis | 146 |

Finally, in Table 4 some virtual optimal conditions for minimal *f*^−^ rate were chosen, i.e. *p*_0_ = 0, sub-linear shape and related unbalanced design. The increase in the superiority parameter over an assumed standard NTP trial is substantial: up to over 200%. An optimal design with minimum *f*^−^ results for the Armitage test, as a scores test of a linear regression in weighted glm, at the steepest slope *b* and the lowest *π*. The steepest increase is obtained for a 2-point design [*D* = 0; *D* = *D*_*max*_] and *n*_*i*_ = *N/*2. The lowest variance is obtained for higher *n*_*i*_ and lower *p*_*i*_. These conditions are not known a-priori and the different demands show a trade-off.

**Table 4:** Optimal condition

| No. | Proportions | Samples sizes | Dose levels | f- | Scenario | ξ-factor |
| --- | --- | --- | --- | --- | --- | --- |
| 1 | 0.01,0.05,0.15,0.2 | 50,50,50,50 | 0,10,50,100 | 0.0702 | base $N^{\text{NTP}}$ | - |
| 2 | 0,0,0,0.20 | 140,20,20,20 | 0,10,20,40 | 0.00049 | $p_0 = 0$, shape sub-linear, opt. $n_0$ | 223 |
| 3 | 0,0,0,0.20 | 140,10,10,40 | 0,10,20,40 | 0.00019 | $p_0 = 0$, shape sub-linear, opt. $n_0$ | 241 |
| 4 | 0,0,0,0.20 | 140,10,10,40 | 0,10,20,1000 | 0.000005 | $p_0 = 0$, shape sub-linear, opt. $n_0$ | 259 |

**Table 5:** Optimal design examples

| No. | Proportions | Samples sizes | Dose levels | f- | Scenario | ratio-to-balanced $\zeta$ |
| --- | --- | --- | --- | --- | --- | --- |
| a | 0.01,0.05,0.15,0.2 | 50,50,50,50 | 0,10,50,100 | 0.0639 | balan. | - |
|  | 0.01,0.05,0.15,0.2 | 110,30,30,30 | 0,10,50,100 | 0.0145 | unbal. | 143 |
| b | 0.00,0.04,0.09,0.19 | 50,50,50,50 | 0,10,50,100 | 0.0409 | balan. | - |
|  | 0.00,0.04,0.09,0.19 | 110,30,30,30 | 0,10,50,100 | 0.0061 | unbal. | 144 |
| c | 0.01,0.01,0.01,0.20 | 50,50,50,50 | 0,10,50,100 | 0.0076 | balan. | - |
|  | 0.01,0.01,0.01,0.20 | 110,30,30,30 | 0,10,50,100 | 0.0092 | unbal. | 97 |
| d | 0.01,0.20,0.20,0.20 | 50,50,50,50 | 0,10,50,100 | 0.4009 | balan. | - |
|  | 0.01,0.20,0.20,0.20 | 110,30,30,30 | 0,10,50,100 | 0.04181 | unbal. | 689 |
| e | 0.01,0.20,0.20,0.15 | 50,50,50,50 | 0,10,50,100 | 0.6713 | balan. | - |
|  | 0.01,0.20,0.20,0.15 | 110,30,30,30 | 0,10,50,100 | 0.1365 | unbal. | 247 |

#### 3.1.1 Optimal design

Already [20] questioned the optimality of the balanced NTP design and proposed an unbalanced design where in control and the high dose most sample sizes are localized. The optimal design for a poly-k trend test or at least the Armitage trend test behind it is quite complex. In the following, the *f*^−^ for (more than dual control) unbalanced designs are empirically compared with balanced designs for different dose-response profiles (additive shift, including *p*_0_ = 0) using a ratio-to-balanced factor *ζ*.

Case a) shows a clear increase of the factor (ie. decrease of *f*^−^) of the unbalanced design. For *p*_0_ = 0 *f*^−^ remains approximately unchanged (case b) compared to *p*_0_ = 0.01. For the sub-linear profile, on the other hand, the unbalanced design shows no change in *f* - with respect of the balanced design (case c), whereas for a plateau shape there is an extreme reduction of *f*^−^ in the unbalanced design (case d). Even for shape with a downturn effect at the high dose an increase of the factor (ie. decrease of *f*^−^) of the unbalanced design occur.

In summary, a partially unbalanced design even more than dual control design almost always appears to have significantly lower f-values than the usual balanced design in the guidelines. This behavior will certainly be different if the estimators of historical controls are used directly in a test statistic (which is probably not yet available for poly-3). Recent work [20, 23] on this also recommends unbalanced designs

In summary, false negative rates for the analysis of crude tumor rates by the Armitage trend test can vary massively depending on data and design, which is not immediately apparent in the routine evaluation of many tumors by toxicologists. Thus, evaluation based on a p-value alone from either crude Armitage or poly-k trend test may not be appropriate for any tumor site, i.e., the current common practice is inappropriate for risk assessment.

### 3.2 False positive rates taking rare events into account

An important *f*^+^ issue (with quite an impact on the *f*^−^ rate as a further consequence) is the exhaustion of the nominal *α*-level of particular tests for these discrete data, already for a single tumor rate alone, much more so for multiple correlated tumor rates [12]. This is another dimension complex for weighted rates in the poly-k approach and furthermore in 2 by k designs. Here, a simple 2 by 2 design is considered as an example, e.g. incidences of hepatoblastoma in female mice for control and the high dose *p*_0_ = 0*/*50; *p*_*max*_ = 11*/*50 from dataset NTP-TR491 [3]. Starting from the asymptotic Pearson test (its Yates-type correction), the Fisher exact test, the conditional tests (asymptotic and mid-p value [39]) [19], the unconditional test and Berger’s [4] p value interval are compared in Table 6.

**Table 6:** One-sided p-values for different tests in 2 by 2 table data

| Test principle | p-value | exhausted size |
| --- | --- | --- |
| Pearson | $2.19 \cdot 10^{-4}$ | 0.0256 |
| Pearson, Yates corrected | $6.97 \cdot 10^{-4}$ | |
| Fisher exact | $2.64 \cdot 10^{-4}$ | |
| Conditional inference, asymptotic | $2.34 \cdot 10^{-4}$ | |
| Conditional inference, mid p value | $1.32 \cdot 10^{-4}$ | |
| Unconditional inference | $1.08 \cdot 10^{-4}$ | |
| Berger's p value interval | $[p_0 < 10^{-10}; p_1 = 2.64 \cdot 10^{-4}]$ | |

The size of about only *α/*2 for the mid-p value approach (available in the CRAN package coin [19]) alone clearly shows the potential for exhausting the nominal *α*-level through suitable tests. Berger’s approach [4] reveals the enormous degree of conservatism by the width of the interval, where *p*_0_ is the smallest p-value achievable if no conservatism occur due to discreteness). I.e., needed is an unconditional approach for 2 by k table data (analogously to [22] and trends [35] [33]) for weighted proportions for a single tumor incidence as well as multiple correlated proportions (see e.g., [30]).

### 3.3 Further influences on the false negative rate of the poly-k test

In chapter 3.1, it will be apparent how sensitive the *f*^−^ rate is to the shape of dose-response relationship (including occurring downturns at high doses) and the dose scores that match it. However, since this is not known a priori, a robust test must be applied. A suggestion is to use a Tukey-type [36] maximum-test over linear, ordinal and logarithmic dose scores instead of linear regression [32]. In addition, to be robust to plateau-shape to effect reversal, dose should be modeled quantitatively and qualitatively (e.g. using Williams contrast [37]) [18]).

Another aspect is the appropriate choice of the Weibull shape parameters k. Usually the *k* = 3 recommendation is followed, sometimes one analyzes with k=3 and 6 in parallel (which increases the f+ rate significantly). By means of the multiple marginal model approach (mmm) [27] one can use all conceivable k-values within a correlated maximum test and thus use the best fitting k-value, of course at the increased conservatism penalty of multiple testing [12].

Two-sided testing massively increases the *f* − rate, so only one-sided hypothesis formulations should be used, since only increasing tumor rates are of interest.

In such a completely randomized design, the risk difference is the natural effect size, even not the canonical link function in the glm-based logistic regression model. In principle, one can construct a maximum test on the three possible effect size, i.e. risk difference, risk ratio or odds ratio [12], [18].

Both in a guideline [2] and in recent publications [29] both trend test and pairwise tests are proposed or used. This cause many decisions, which may be contradictory, and increase further the false positive rate [25]. One can speculate about the reasons for such a complex approach, but certainly the availability of the individual inference of each dose *D*_*i*_ vs. C-plays a role (which a global trend test does not provide) which is especially robust to downturn effects at high doses. This is achieved by a joint Williams and Dunnett-type test [21] or Tukey-type and Dunnett-type test [18] that controls the FWER at the cost of further increased conservatism (i.e., increase in *f*^−^rate), whereas poly-k adjustment is available for both approaches [16].

Ultimately, the question remains whether a significant p-value of a poly-3 trend test for at least one tumor site in male or female animals is sufficient for the global conclusion that the study is positive (and if not, what other criteria to use). In the sense of reducing complexity, there is a great desire for a dichotomous evaluation (+ or -), whereby a third category, i.e. undecidable, presumably concerning the majority of the studies, would be important - but is not explicitly demanded.

The sex-specified evaluation, each at level *α*, represents the standard evaluation. This is without alternative for specific tumors, such as leydig cell carcinoma. However, the vast majority of tumors occur in both sexes, which can then be analyzed together appropriately [13], which reduced data-dependent the false negative rate seriously by up to doubling *n*_*i*_.

According to the guidelines, 4 independent studies are usually performed within a bioassay, two-species-two-sex study (i.e. mice/rat/males/females). For glyphosate, the data from 10 long-term bioassays are available for males and females, i.e. 20 data sets. What influence do 20 data sets have on the final statement compared to 4 according to the guideline? On the one hand, the total sample size *N*_*total*_ increases, on the other hand, there is a much greater between-assay variability, and in addition, the dose ranges of the studies are very heterogeneous. A joint related mixed effect model may help to overcome this difficulty, see e.g. [17].

Historical controls can be helpful, either indirectly through post-testing interpretation using their prediction intervals [26] or direct inclusion in the test statistics; for the poly-3 test alone, no statistical procedure appears to be described, nor does any software exist. Furthermore, this requires a new quality of historical controls, since not only tumor-specific incidences but animal-specific mortalities must be available to make the poly-k estimators available.

## 4 Proof of hazard vs. proof of safety

The central argument in [31] *‘primary difficulty with animal bioassay data is not high false positive rates, but low statistical power*’ can be rephrased into *f*^−^ is the more important error rate (when *f*^−^ = 1 − *power*), one should control it directly, since the Neyman-Pearson hypothesis system is asymmetric and controls only one error rate directly, just the more important, just ^−^ here. This leads to the proof of safety, already available for decades [6, 10], but hardly used nowadays, see e.g. aquatic bioassays [9]. One would have to use non-inferiority tests, which necessarily require the a priori definition of a still tolerable tumor rate, which should be site and tumor-specific, at least probably specific for adenomas/carcinomas or rare/common tumors. This alone is challenging, in addition there would be frighteningly high required *n*_*i*_, or for *n*_*i*_ = 50 the insight how little, and this extremely tumor-specific, quantitative ‘safety’ would be detectable (or the other way around, how high *f*^−^’s one would have to accept). The proof of safety demonstrate non-inferiority of all *q* tumors using an intersection-union test (IUT) based on *q* marginal level-*α* tests [11]. An IUT in this dimension *q* is extreme conservative ([14]) that a global proof is unlikely to succeed on the basis of real data. If one adds the safety demonstration by *k* marginal level-*α* non-inferiority tests between each dose and negative control, then the dimension increases even to *q * k*. On the other hand, from the perspective of a pronounced proof of hazard, one would consider the *q* tumors as primary multiple endpoints by demonstrating union-intersection test (UIT) based on *q* correlated tests. An UIT of this dimension *q* is also conservative especially since the correlation of so many multiple binaries is not too large that one could substantially mitigate the conservative Bonferroni level *α/q*. The conservatism of these discrete tests can be mitigated somewhat by maximal exhaustion of the nominal significance level [30] (using the CRAN library *multfisher* for a 2-sample design with correlated binary endpoints, see e.g. [15].

Thus, the current approach of interpreting independent marginal level *α*-tests is in itself methodologically trivial but inherently associated with a massive increase in the false positive rate: this would practically approach 100% for the common *>* 50 tumor entities.

It is easy to see that all 3 approaches are extremely different with respect to the underlying model and thus come to very different conclusions of toxicological risk assessment, which study should be considered positive (ie. at least a significant p-value). The guidelines do not explicitly address this important problem and a consensus on a recommended test version would be needed. Until then, none of the three is appropriate or inappropriate per se.

## 5 Summary

Numerous data and design-dependent effects massively influence the *f*^+^*/f*^−^ ratio of the poly-k test. This is rarely obvious, but influences the interpretation considerably. Particularly, the influence of multiple correlated tumors analysis, the shape of dose-response relationship depending on the design (as a result of the study) were discussed.

## Acknowledgment

^1^ My special thanks to Dr. A. Szabo, Medical College of Wisconsin, for providing R code for various unpublished methods of power calculation of the Armitage trend test.

